# Annotation-Informed Causal Mixture Modeling (AI-MiXeR) reveals phenotype-specific differences in polygenicity and effect size distribution across functional annotation categories

**DOI:** 10.1101/772202

**Authors:** Alexey A. Shadrin, Oleksandr Frei, Olav B. Smeland, Francesco Bettella, Kevin S. O’Connell, Osman Gani, Shahram Bahrami, Tea K. E. Uggen, Srdjan Djurovic, Dominic Holland, Ole A. Andreassen, Anders M. Dale

**Affiliations:** NORMENT, Institute of Clinical Medicine, University of Oslo and Division of Mental Health and Addiction, Oslo University Hospital, 0407 Oslo, Norway; Department of Medical Genetics, Oslo University Hospital, Oslo, Norway; NORMENT, Department of Clinical Science, University of Bergen, Bergen, Norway; Multimodal Imaging Laboratory, University of California, San Diego, La Jolla, CA, USA; Department of Neurosciences, University of California, San Diego, La Jolla, CA, USA; Department of Radiology, University of California, San Diego, La Jolla, CA, USA; Department of Psychiatry, University of California, San Diego, La Jolla, CA 92093, USA

**Keywords:** genetic architecture, complex traits, functional genetic categories, polygenicity, variant effect size

## Abstract

Determining the contribution of functional genetic categories is fundamental to understanding the genetic etiology of complex human traits and diseases. Here we present Annotation Informed MiXeR: a likelihood-based method to estimate the number of variants influencing a phenotype and their effect sizes across different functional annotation categories of the genome using summary statistics from genome-wide association studies. Applying the model to 11 complex phenotypes suggests diverse patterns of functional category-specific genetic architectures across human diseases and traits.

## Background

The rapid technological advances of the last years have provided an enormous amount of genetic data thereby promoting the development of statistical methods aimed to unravel the genetic architecture of complex traits [1]. A key effort has been to estimate SNP-based heritability, either using individual-level genotype data [2], or summary-level statistics from genome-wide association studies (GWAS) [3]. However, heritability estimates provide a limited picture of the genetic architecture underlying complex phenotypes. For example, they are agnostic about the number of genetic variants influencing a phenotype and their effect sizes [4]. Both of these quantities can vary and still result in the same heritability, which is proportional to their product [5, 6].

Recently, we developed a model which allows the breakdown of SNP-heritability into the number of variants influencing a given phenotype (non-null variants) and the distribution of their effect sizes using summary statistics from GWAS and detailed population-specific linkage disequilibrium (LD) structure [5, 6]. This model assumes a uniform distribution of non-null variants with the same expected effect size throughout the genome. However, prior genetic studies suggest that non-null variants are differentially enriched across functional genomic categories and complex phenotypes [7, 8]. Here we present a model, Annotation Informed (AI) MiXeR, which extends our previous work by allowing different (non-overlapping) predefined functional annotation categories of the genome to have various densities of non-null variants with different effect size distributions.

Several conceptually related methods aiming to characterize the genetic architecture of phenotypes using GWAS summary statistics have recently been developed. The partitioned LD score regression (LDSC) analysis presented by Finucane and colleagues [9] estimates the proportion of SNP-based heritability explained by variants within predefined functional categories but does not estimate the abundance of non-null variants or assess their effect sizes. The RSS-E method [10] only estimates the abundance of non-null variants in different annotation categories, while the distribution of effect sizes of non-null variants is assumed to be the same throughout the genome. Alternatively, the GENESIS model [11] allows several groups of trait-susceptibility variants with different densities and effect size distributions but assumes the non-null variants to be uniformly distributed among the groups and does not support prior group definition (e.g. in terms of functional annotation categories). AI-MiXeR allows simultaneous modeling of abundance and effect size magnitudes of non-null variants in arbitrary predefined functional annotation categories.

We extensively tested AI-MiXeR on synthetic GWAS data generated under various setups to establish scenarios where it reconstructs the underlying parameters correctly. We then applied AI-MiXeR to GWAS summary statistics for 11 complex phenotypes representing a range of diverse human traits and diseases. This analysis suggests that both densities and effect sizes of non-null variants can vary considerably in different genomic annotation categories and reveals diverse patterns of genetic architecture in different phenotypes.

## Results

### AI-MiXeR model overview

We consider an additive model of genetic effects ignoring gene-environment interactions, epistasis and dominance effects. Variant effect sizes are modeled with point-normal mixture priors, where both proportion of non-null variants and distribution of their effect sizes can vary between different predefined functional genomic categories. Each functional category in the model is characterized by the proportion of non-null variants (polygenicity, π) and the variance of their effect sizes (discoverability, σ^2^). The pure (i.e. not induced by LD) effect of the k^th^ variant (*β*_*k*_) is modeled as a mixture of null and causal (see Methods) components: 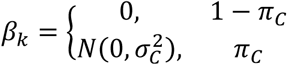, where *π*_*c*_ and 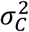 respectively are proportion and variance of non-null variants effect sizes in the functional category *C*. The signed association test statistics (z-score) of the j^th^ variant is then given by: 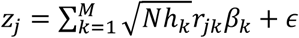, where *N* is the GWAS sample size, *h*_*k*_ is the heterozygosity of variant *k, M* is the number of variants in LD with variant *k, r*_*jk*_ is the Pearson’s correlation coefficient between the genotypes of the j^th^ and k^th^ variants and *ϵ* is a 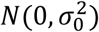 distributed residual factor. Functional category-specific polygenicities and discoverabilities of a GWAS trait are estimated by maximizing the likelihood of the GWAS summary statistics (z-scores). To reduce computational burden, we randomly select a subset of 10^6^ variants of all GWAS variants to use for maximization of the likelihood function. For specific details of the model and its implementation please refer to the Methods section.

### Simulations with synthetic data

We analyzed the performance of the model on GWAS summary statistics generated from synthetic genotypes and phenotypes with various genetic architectures under model assumptions.

#### Synthetic genotypes

10^5^ synthetic genotypes were generated with Hapgen2 [12] using 503 European samples from 1000 Genomes Phase 3 data [13] as described in [6]. A set of 11,015,833 biallelic variants was considered. The LD structure was estimated from a subset of 10^4^ genotypes using PLINK 1.9 [14] ignoring all correlations between variant genotypes at *r*^2^ < 0.01 and trans-chromosome correlations.

#### Functional annotation categories

Two non-overlapping functional annotation categories were considered: exonic and non-exonic. The exonic annotation category includes all variants within exons (including 5’ and 3’ untranslated regions) of protein coding genes, while the non-exonic category contains all remaining variants. This choice was motivated by previous research showing that protein coding exons (including 5’ and 3’ untranslated regions) are most strongly enriched of association with many complex human phenotypes [8]. Additionally, the exonic category, as defined above, largely overlaps with the genomic regions investigated in whole exome genotyping and whole exome sequencing studies. Its modeling can therefore serve as a projection for future discoveries in whole exome studies. All variants were functionally annotated using UCSC’s Table Browser (hg19/GRCh37) [15]. With this definition, the non-exonic category contains approximately 70 times more variants than the exonic category.

#### Synthetic phenotypes

were generated using SIMU [16]. A given number of non-null variants was selected at random for each functional annotation category. Effect sizes for the selected non-null variants were sampled from the standard normal distribution and then rescaled to obtain the required level of heritability, given different predefined ratios (see below) between the average effect sizes of the two dichotomous functional annotation categories. For each synthetic genotype, a quantitative synthetic phenotype was then generated as the sum of allelic effects over all non-null variants complemented by a certain proportion of a random Gaussian noise (representing effects of the environment) required to keep the predefined level of heritability. Finally, association tests were performed using PLINK 1.9 to obtain GWAS summary statistics.

#### Simulation setup

All possible combinations of the following parameter values were used for generating synthetic phenotypes: 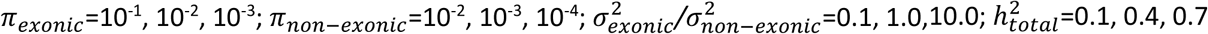, resulting in 81 different parameter setups covering a broad range of genetic architectures. Ten different phenotypes with independently generated locations of non-null variants and effect sizes thereof were generated for each combination of parameters, resulting in 810 synthetic phenotypes (and corresponding GWAS summary statistics).

The simulations demonstrate that the true parameters are estimated accurately when the proportions of heritability carried by both functional categories are comparable and each category individually carries >2% of the total heritability; if one of the functional categories carries a negligible fraction (<2%) of the total heritability the model often fails to reconstruct its parameters accurately (Supplementary Figure 1). Selected simulation cases representing scenarios closely resembling phenotypes analyzed in this study are presented in Figure 1. These simulations show that in the range of parameters observed (according to the model) in the 11 phenotypes analyzed in this study, the model is able to provide instructive unbiased estimates of *π* and *σ*^2^ parameters for both exonic and non-exonic functional annotation categories. A complete comparison of true simulation parameters and corresponding model estimates for all 810 simulated phenotypes is shown in Supplementary Figures 2 – 4, and the corresponding numerical results are given in Supplementary Table 3. Of note is that, in general, heritability estimates are more robust than estimates of *π* and *σ*^2^.

**Figure 1.**
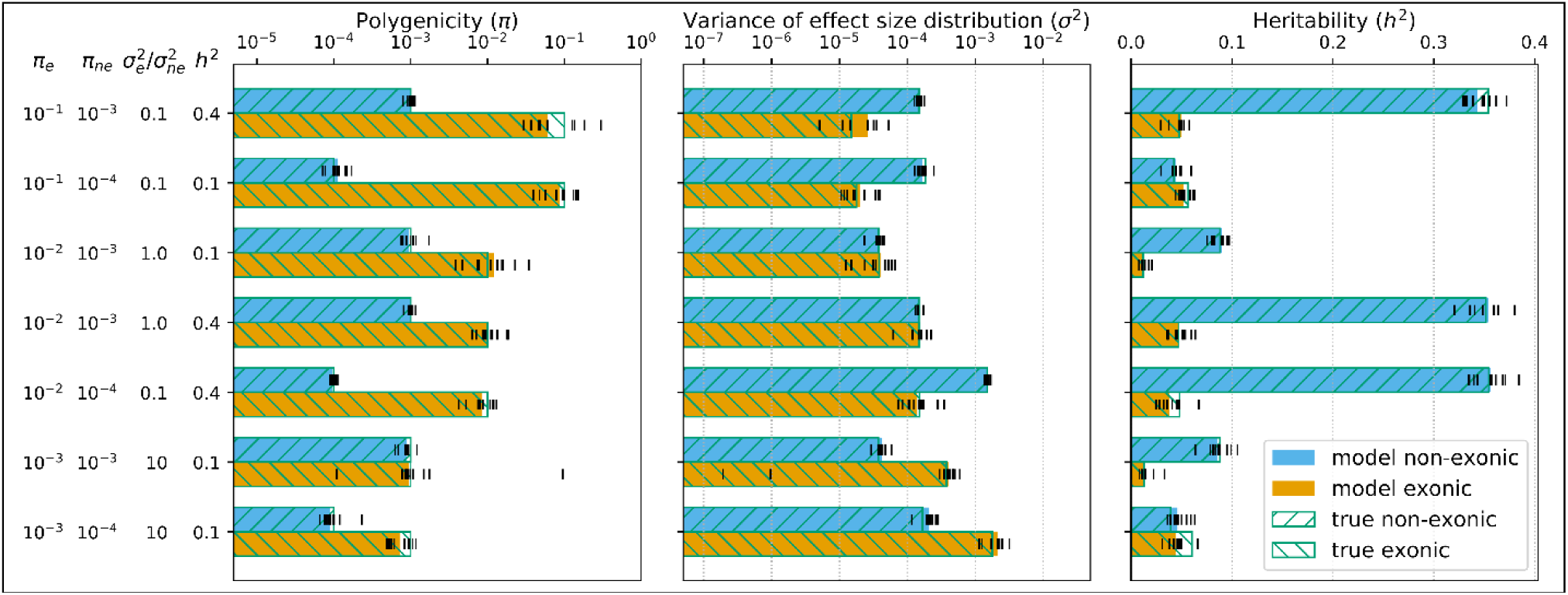
Performance of the model on a selected set of scenarios with synthetic GWAS data. True simulation parameters (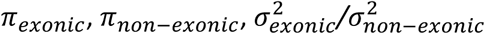 and 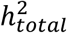) are shown on the left. The blue bars represent the non-exonic category, the orange bars represent the exonic category. The bar lengths represent the median values obtained from 10 optimization runs with independently generated GWAS (locations and effect sizes of non-null variants). The parameter values from individual optimization runs are shown with vertical black dashes. Empty bars with green borders and hatches show the true values of the corresponding parameters used for GWAS simulation.

### GWAS summary statistics

We applied the model to GWAS summary statistics on 11 phenotypes (Figure 2, Supplementary Table 1) including schizophrenia (SCZ) [17], bipolar disorder (BD) [18], attention deficit/hyperactivity disorder (ADHD) [19], general cognitive ability (COG) [20], educational attainment (EA) [21], type 2 diabetes (T2D) [22], inflammatory bowel disease (IBD) [23], low-density lipoproteins (LDL) [24], body mass index (BMI) [25], height [25] and waist-hip ratio (WHR) [26]. Details on these studies including year of publication and number of samples (cases/controls when appropriate) are given in Supplementary Table 4.

**Figure 2.**
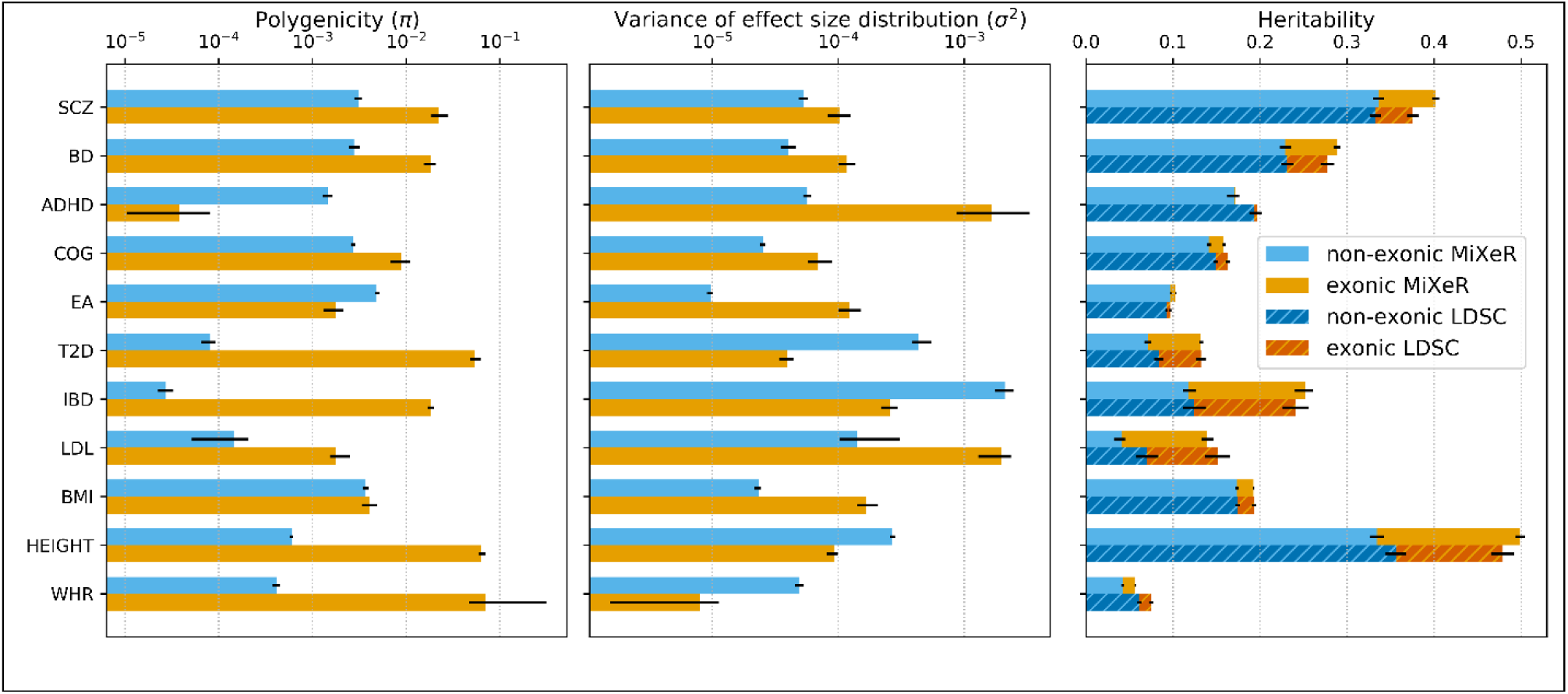
Estimated polygenicity, discoverability and heritability of exonic and non-exonic functional categories in 11 traits. Traits are shown in the left column: schizophrenia (SCZ), bipolar disorder (BD), attention deficit/hyperactivity disorder (ADHD), general cognitive ability (COG), educational attainment (EA), type 2 diabetes (T2D), inflammatory bowel disease (IBD), low-density lipoproteins (LDL), body mass index (BMI), height and waist-hip ratio (WHR). For AI-MiXeR results, a bar’s length shows the mean value of the parameter obtained from 50 independent optimization runs with 10^6^ randomly selected variants used to maximize the likelihood of the observed GWAS z-scores, while the black bars show the range (min, max) of such estimates. The heritability estimates obtained with partitioned LDSC are shown in darker colors with hatching (right) and black bars representing the standard errors of the estimates.

Like in simulations with synthetic data, we consider here two functional annotation categories for the variants: exonic and non-exonic. For ease of comparison with partitioned LDSC method [9], the LD structure was estimated with PLINK 1.9 using genotype data from LDSC’s template [27] containing 9,997,231 biallelic variants for 489 unrelated European individuals (originally derived from 1000 Genomes Phase 3 data [13]). Trans-chromosome correlations as well as correlations between variant genotypes at *r*^2^ < 0.05 were disregarded. We also obtained estimates of SNP-heritability per functional category for all 11 phenotypes using partitioned LDSC. These estimates are compared with AI-MiXeR’s in Figure 2 (right) and Supplementary Table 1. The polygenicity parameters (π_exonic_ and π_non-exonic_) can be converted into the number of non-null variants by multiplying them by the total number of variants within the corresponding annotation category. The numbers ensuing for the analyzed phenotypes are presented in Supplementary Table 1. For each phenotype, 50 independent optimization runs were performed to maximize the likelihood of the GWAS z-scores observed in different subsets of 10^6^ randomly selected variants.

The majority of analyzed phenotypes fall into the portion of parameter space where, according to our simulations, the model is expected to produce robust parameter estimates (Supplementary Figure 1, Figure 1). However, two phenotypes (ADHD and WHR) fall in a portion of parameter space where the model is prone to return inconsistent results in the exonic category (due to this category’s limited size, its very low polygenicity in ADHD and its very low discoverability in WHR). This is reflected in larger error bars for the exonic category in these phenotypes (Figure 2, Supplementary Table 1). However, the observed consistency of parameter estimates across all 50 independent optimization runs for all analyzed phenotypes encourages to draw conclusions about actual features of the underlying genetic architecture.

## Discussion

We present the AI-MiXeR model, which can be used to decouple and partition a phenotype’s heritability into functional category-specific polygenicity (proportion of non-null variants) and discoverability (variance of non-null effect sizes) components and thus better characterize the phenotype’s genetic architecture.

It is widely assumed that protein coding exons contain a higher proportion of causal variants (higher polygenicity) and have on average stronger effects (higher discoverability) on complex phenotypes when compared to non-exonic regions [8, 28, 29]. In our study, the AI-MiXeR model suggests that less than half (5 of 11) of the analyzed phenotypes (SCZ, BD, COG, LDL and BMI) support this assumption. Four other phenotypes (T2D, IBD, height and WHR) show higher density of non-null variants in exonic regions but stronger average effects in the non-exonic portion of the genome. In the two remaining traits (ADHD and EA), the pattern appears to be reversed, with a higher density of weaker effect variants in non-exonic regions.

According to the model, EA has the largest number of non-null variants (48000, with only 0.6% of exonic variants) among all analyzed phenotypes, while IBD has the smallest (3000, 91% exonic) (Supplementary Table 1). However, the effects of the non-null variants are on average 5 times stronger in IBD than in EA and result in a larger total heritability for the former (0.25 in IBD vs 0.1 in EA). SCZ and BD show similar polygenicity (34000 and 30000 non-null variants, of which 9% and 10% respectively are exonic) and discoverability (with exonic effects being approximately twice stronger). In contrast, most non-null variants for height and T2D are exonic (62% out of 15500 and 90% out of 9000 respectively) having on average 10- and 3-times weaker effects respectively than non-exonic variants.

The model suggests, that despite having similar heritability, phenotypes may differ substantially in polygenicity and discoverability of non-null variants. For example, both AI-MiXeR and partitioned LDSC provide comparable estimates of total and partitioned heritability for LDL and T2D (AI-MiXeR LDL: 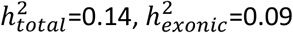 AI-MiXeR T2D: 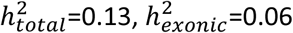) (Figure 2, Supplementary Table 1, Supplementary Table 2). However, our model suggests that the genetic architectures underlying these phenotypes differ drastically, with T2D being approximately 5 times more polygenic than LDL and having 91% (vs 15% in LDL) of non-null variants within exons. The polygenicity deficit is compensated in LDL with a discoverability 3.5 times larger than in T2D (50 times larger in the exonic category).

From our simulation studies on synthetic GWAS, we can infer that the balance of *h*^2^ partition between the functional annotation categories has a strong effect on the model’s performance. Extremely small values of polygenicity (*π*) or discoverability (*σ*^2^) in a functional annotation category (relative to the complementary category) result in a heavily unbalanced heritability partition between the categories and can thus lead to substantial errors in the estimates of *π* and *σ*^2^ for the category with smaller absolute heritability (Supplementary Figure 1 top and bottom). Despite this, heritability estimates were generally robust (Supplementary Figure 1 – 4, Supplementary Table 3).

Decoupling the heritability of different functional categories into polygenicity and discoverability may facilitate trait-specific experimental designs prioritizing certain genomic regions for detailed investigation. For instance, by looking only at the heritability pertaining exons in T2D 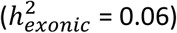 and LDL 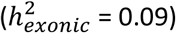, one could expect the yield of an exome-wide scan for both phenotypes to be comparable. However, AI-MiXeR predicts that the average effect size (square root of discoverability) of exonic non-null variants is approximately 7 times larger in LDL than in T2D. An exome study of the former could be therefore expected to result in a higher yield of statistically significant findings, given a moderately sized sample. This speculation may be indirectly supported by comparing existing exome-wide studies of T2D and LDL. One of the largest exome sequencing studies on T2D (20791 cases, 24440 controls) published so far identified 15 variants in 7 distinct genomic loci reaching exome-wide significance level [30]. In contrast, an exome-wide association study of serum lipids in a comparable sample (N = 39087) reported 66 exome-wide significant LDL susceptibility variants within 14 loci [31]. AI-MiXeR’s predictions also suggest, however, that a significant increase in the sample size for T2D will yield more fruitful results than an equivalent sample size increase in LDL, since T2D has substantially larger polygenicity.

AI-MiXeR relies on design and implementation quality of the specific GWAS. In general, model estimates for the same phenotype may differ depending on a GWAS’s sample size, as well as on the coverage of the tested variants. The samples of the GWAS tested here vary by more than one order of magnitude in size, from approximately 5×10^4^ for BD and ADHD to more than 7×10^5^ for EA and height. In all simulations, we kept the sample size constant (N = 10^5^) and varied only the heritability (*h*^2^=0.1, 0.4, 0.7). Since these quantities contribute to the GWAS z-scores distribution only through their product (follows from formula (5)), our simulation scenario with N=10^5^ and *h*^2^=0.7 is equivalent to a scenario with, for example, N=7×10^5^ and *h*^2^=0.1. Other aspects of potential GWAS-related issues (e.g. coverage of tested variants) were not tested.

The model underlying AI-MiXeR is sensitive to the LD structure estimates. Ideally, the LD structure should be estimated on the same sample used for association testing. However, this is mostly impractical if not impossible. Here, in the analysis of GWAS summary statistics for 11 phenotypes we estimated the LD structure using the 1000 Genomes Phase 3 genotype panel. Inconsistencies between the LD structure of the samples used for association testing and that of the 1000 Genomes Phase 3 panel could skew the model’s results. Additionally, roughening the LD structure (e.g. by ignoring all correlations with *r*^2^ below a certain threshold) also could result in biased parameter estimates. In our simulations, *r*^2^ was estimated from 10,000 synthetic genotypes (randomly sampled from the complete set of 100,000 synthetic genotypes used for association testing) ignoring all correlations with *r*^2^ < 0.01. A subset of European ancestry samples from the 1000 Genomes Phase 3 panel (N = 489) was used to estimate LD *r*^2^ values for the GWAS data because of the wide availability of these data, ease of comparison with LDSC and the fact that genotypes in majority of analyzed GWAS were imputed using this panel as a reference. The limited size of the 1000 Genomes panel, however, results in relatively low confidence *r*^2^ values, especially for weak correlations involving low-frequency variants. To mitigate this issue, we increased the *r*^2^ cutoff, disregarding all correlations with *r*^2^<0.05. Nevertheless, consistency between partitioned heritability estimates produced by AI-MiXeR and LDSC (Figure 2, Supplementary Table 2) suggests the absence of the model-specific systematic biases.

AI-MiXeR makes further simplifying assumptions, including uniform distribution of non-null variants within functional annotation categories and the effect size’s independence of allele frequency and LD. It has recently been shown that these simplified assumptions, which have been used implicitly or explicitly in many earlier methods, can lead to substantial biases in heritability estimates [32]. We previously demonstrated [6] that these factors also bias the model’s estimates of *π* and *σ*^2^ when no distinction is made between annotation categories. We did not investigate how disregarding them affect AI-MiXeR’s category-specific estimates. These assumptions are likely violated to different degrees in different phenotypes and make the model more suitable for some phenotypes than for others. In the current report, we provide examples of successful applications of AI-MiXeR, but advise prospective users to carefully assess the model’s suitability for a given phenotype.

It is also important to note that the numbers of non-null variants presented (Supplementary Table 1) should be taken with a grain of salt because of the model’s assumptions and technical simplifications. The observed relative proportions of non-null variants (and their effect sizes) between functional annotation categories within one trait or across different traits might be more reliable indicators of actual genetic architecture features or differences.

## Conclusions

The AI-MiXeR method presented here considers predefined annotation categories allowing both different proportions of non-null variants and different effect size distributions in various functional annotation categories, which is not possible with other methods available to date [9-11]. The ability to model predefined annotation categories separately allows hypothesis-driven studies of complex phenotypes, which in turn provide a better understanding of the genetic architecture of those complex phenotypes. Our analysis suggests that both the polygenicity and the discoverability in different functional categories vary considerably across human traits and disorders. Knowing such patterns may facilitate trait-specific experimental designs prioritizing specific genomic regions for detailed investigation.

## Methods

### AI-MiXeR model

Consider a quantitative phenotype standardized to mean 0 and variance 1. Let *y* be a random variable representing a phenotype measurement for an individual in the population (E(*y*) = 0, var(*y*) = 1). Let *G* = {*g*_*j*_}_*j*=1…*M*_ be a fixed set of *M* random variables representing genotypes of bi-allelic variants. These are assumed to be centered (E(*g*_*j*_) = 0) but not scaled (var(*g*_*j*_) = 2*f*_*j*_(1 − *f*_*j*_) = *H*_*j*_, where *f*_*j*_ is the minor allele frequency of variant *j* and *h*_*j*_ is its heterozygosity). We assume an additive genetic model for the phenotype generation:

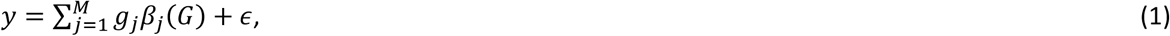

where *ϵ* is a normally distributed error term with mean 0 and variance 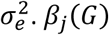 is understood here as the (unknown) effect of variant *j* as would be obtained from a multiple linear regression of the phenotype *y* on all genotypes *G* in a hypothetical infinite sample. This definition of the effect size *β*_*j*_(*G*) implies that *β*_*j*_(*G*) will reflect only the true causal effect of the j^th^ variant (thus *β*_*j*_(*G*) = 0 if the j^th^ variant is not causal) whenever *G* includes all causal variants for the trait. On the other hand, if any causal variants are missing in the set *G, β*_*j*_(*G*) will also include the effects of those missing causal variants that happen to be tagged by the j^th^ variant. Any variant *j* with *β*_*j*_(*G*) ≠ 0 will be called a non-null variant. Further, we will henceforth consider *G* to be fixed and omit it from the notation.

Consider now a GWAS on a quantitative phenotype. Let *N* be the number of individuals in the GWAS and assume that *N* is sufficiently large, so that the allelic composition (i.e. genotype frequencies) of the variants observed in the GWAS is approximately equivalent to the allelic composition of the same variants in the population. Then *ŷ* = (*ŷ*_1_ … *ŷ*_*N*_) is a vector of phenotypes, *Ĝ* = [*ĝ*_*ij*_] _*i*=1…*N,j*=1…*M*_ is the *N* × *M* matrix of genotypes observed in the GWAS and 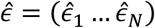 is a vector of residuals (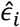 represents the residual term for the i^th^ individual). Using (1) and this notation we can write:

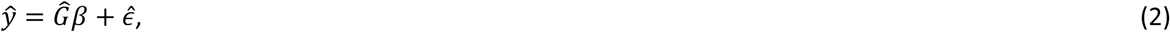

which is a sample-equivalent of (1). Denote also with *ĝ*_*j*_ = (*ĝ*_1*j*_ … *ĝ*_*Nj*_) the vector of genotypes of variant *j* observed in the GWAS (j^th^ column of *Ĝ* matrix). The marginal effect of variant *j* estimated in GWAS 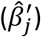 is obtained from the simple linear regression of the phenotype on the genotype of variant 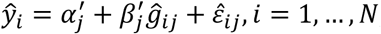, where 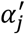 and 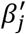 are the (unknown) intercept and slope of the simple linear regression, respectively, and 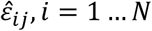, are its residuals. The value of the slope 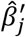 minimizing the sum of squared residuals is:

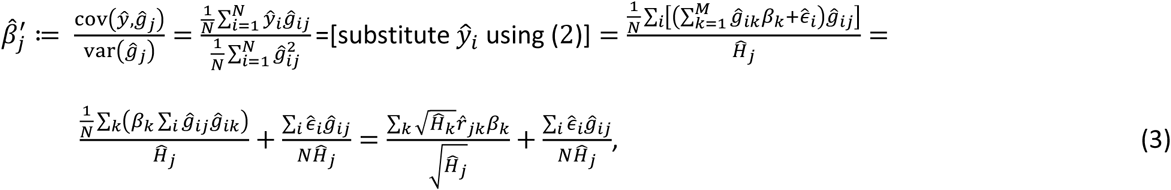

where 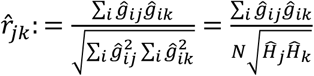 is the sample correlation coefficient between genotype vectors *ĝ*_*j*_ and 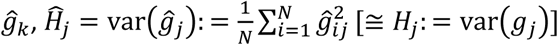 is the sample heterozygosity of variant *j* and *β*_*k*_ is the hypothetical effect size of variant *k* from a multiple linear regression in an infinite population (as discussed above).

Assuming the considered phenotype is complex, i.e. it is influenced by many variants each explaining only a tiny fraction of phenotypic variance, then the variance of the simple linear regression error is approximately equal to the sample variance of the phenotype, 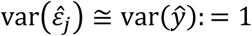, where 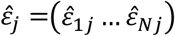. Using this approximation, we can write an expression for the standard error of 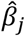:

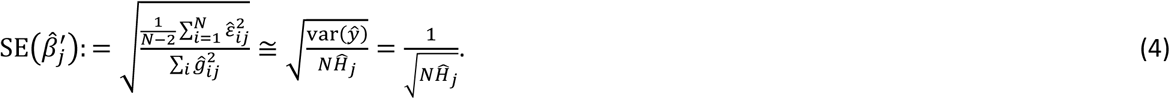

Combining equations (3) and (4) we can write an expression for the z-score of variant *j* observed in GWAS:

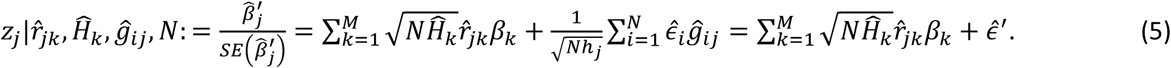

In (5), 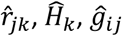 and *N* are known constant factors and 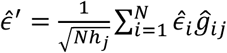 is an unknown (because the 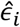 are) residual. Remembering that, by definition, 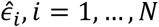 are realizations of the *ϵ* random variable (see equations (1) and (2)) and assuming that these realizations are independent from each other, 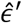 can be modeled as a normally distributed random variable with mean 0 and variance (equally for all variants):

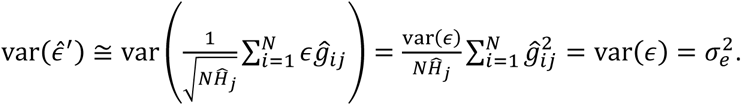

By construction (equation (1)), when there is no genetic effect on the phenotype, 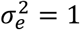. However, the assumption of independence of all 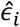 is often violated in GWAS due to the presence of various confounding factors such as population stratification and cryptic relatedness. Moreover, both 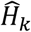 and 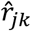 are usually estimated from external genotyping panels where variant frequencies and correlations may differ from those in the GWAS sample. In addition, due to technical limitations, 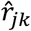 estimates are commonly truncated (e.g. disregarding all correlations below a certain threshold). For the model to be able to mitigate these discrepancies we introduce a 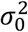 parameter and model 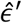 in (5) as a random variable distributed as 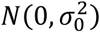. It was shown that, in the framework of the infinitesimal model, 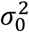 has the same mathematical meaning as the intercept term in the LDSC model [6]. The last unknown factor in (5), *β*_*k*_, is modeled as a random variable with point-normal mixture distribution, where the variance is allowed to differ between different variant annotation categories:

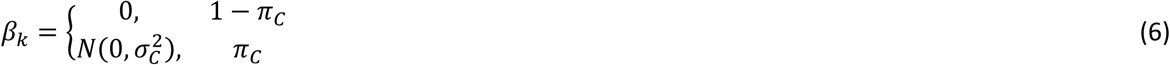

where variant *k* ∈ *C, C* ⊆ *G* is a subset of variants in *G* constituting some annotation category, *π*_*C*_ is the proportion of variants with non-zero effect (non-null variants) in the annotation category *C* and 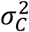 is the variance of the effect sizes among all non-null variants in *C*. The set of annotation categories {*C*_*j*_}_*j*=1…*T*_ defined on *G* must form a partition of *G* (i.e. each variant from the *G* must belong to one and only one annotation category *C*_*j*_).

Modelling *β*_*k*_ as (6), 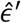 as 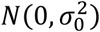 and taking *r*_*jk*_, *h*_*k*_ and *N* as known constant factors, (5) allows to derive the probability density function of *z*_*j*_ (*pdf*_*z*_) as the convolution of *β*_*k*_ (*k* = 1 … *M*) and 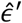 random variables.

### Estimation of probability density function of z-scores

We derive the probability density function (pdf) of a random variable *z* representing a variant’s association z-score from the convolution of *β*_*k*_ (*k* = 1 … *M*) and 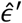 random variables. To simplify notation, we omit the indices reflecting the annotation category, replace 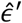 with *ϵ* and denote:

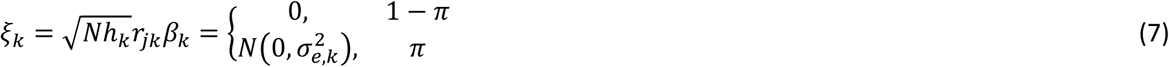

where 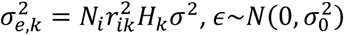. We can then rewrite (5) as:

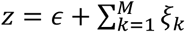

The probability density function of *z* at *z*_0_ (in our case *z*_0_ is the z-score from the GWAS) can be written as the inverse Fourier transform of its characteristic function *ϕ*_*z*_(*t*):

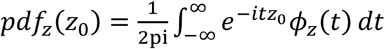, where pi is Archimedes’ constant (i.e. pi ≈ 3.14) and *i* is the unit imaginary number.

Assuming that the non-null effects (*β*_*j*_) are independent from each other and from the error term *ϵ*:

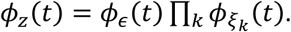

Using the definition of characteristic function and expression (7), we can write the characteristic function of *ξ*_*k*_ as:

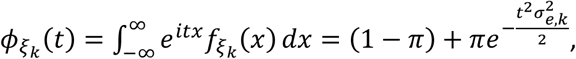

and similarly for *ϵ*:

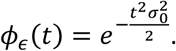

Combining the last two expressions, the characteristic function of *z* can be written as:

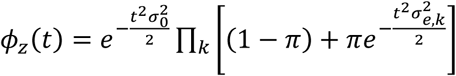, from which we can obtain the point estimate of *pdf*_*z*_at *z*_0_:

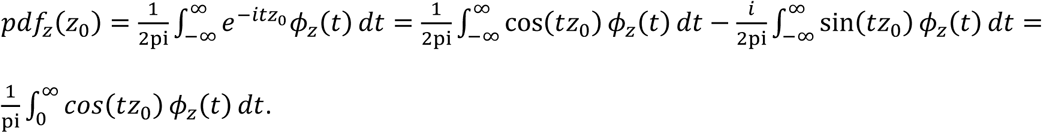

The result is a definite integral (i.e. a number), which can be computed numerically.

### Optimization setup

The polygenicity (*π*) and discoverability (*σ*^2^) parameters are estimated by maximizing the likelihood of the z-scores observed in the GWAS summary-level data 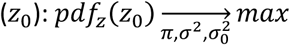, where the probability density function of the z-scores (*pdf*_*z*_) is modeled as described in the section above. Specific estimation details are given below.

The following optimization setup was used:

- Nelder-Mead method (maxiter=1200, fatol=1e-7, xatol=1e-4, adaptive=True) was applied starting from the best point obtained after a single iteration of differential evolution (maxiter=1, popsize=50, init=latinhypercube, bounds: 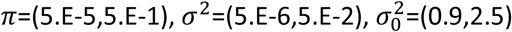). SciPy [33] implementations of both Nelder-Mead and differential evolution methods were used.
- Variants from the extended major histocompatibility complex (MHC) region (genome build 19 locations chr6:25119106–33854733) were excluded from the optimization due to the high complexity of the LD structure in this region.
- The z-scores of 10^6^ randomly selected variants were used for the optimization of the cost function. This procedure was replicated 50 times to limit selection bias.
- The cost function was defined as -log(likelihood)/10^6^, where 10^6^ reflects the number of variants (z-scores) used at each replica of the optimization procedure.

## Declarations

### Ethics approval and consent to participate

Not applicable

### Consent for publication

Not applicable

### Availability of data and materials

The AI-MiXeR method described in this paper is available as a part of the MiXeR repository on GitHub [34]. LD Score Regression software: https://github.com/bulik/ldsc. 1000 Genomes Project Phase 3 variant calls and population data: ftp://ftp.1000genomes.ebi.ac.uk/vol1/ftp/release/20130502/. Psychiatric Genomics Consortium GWAS summary statistics data on SCZ, BD and ADHD: https://www.med.unc.edu/pgc/results-and-downloads. CTGlab GWAS summary statistics data on COG: https://ctg.cncr.nl/software/summary_statistics. Social Science Genetic Association Consortium GWAS summary statistics data on EA: https://www.thessgac.org/data. Diabetes Genetics Replication And Meta-analysis consortium GWAS summary statistics data on T2D: https://diagram-consortium.org/downloads.html. International Inflammatory Bowel Disease Genetics Consortium GWAS summary statistics data on IBD: ftp://ftp.sanger.ac.uk/pub/project/humgen/summary_statistics/human/2016-11-07. Global Lipids Genetics Consortium GWAS summary statistics data on LDL: https://csg.sph.umich.edu/willer/public/lipids2013. Genetic Investigation of Anthropometric Traits consortium GWAS summary statistics data on BMI, height and WHR: https://portals.broadinstitute.org/collaboration/giant/index.php/Main_Page.

### Authors’ contributions

AMD, OAA, OF and AAS conceived the research. OF, DH and AAS implemented and tested the method. OAA secured the funding. OBS, OAA and AAS drafted the manuscript. All authors discussed the results, revised the draft and approved the final manuscript.

## Acknowledgements

This work was performed on the Abel Cluster, owned by the University of Oslo and Uninett/Sigma2, and operated by the Department for Research Computing at USIT, the University of Oslo IT-department (http://www.hpc.uio.no/).

## References

1. Evans, L.M., et al., Comparison of methods that use whole genome data to estimate the heritability and genetic architecture of complex traits. Nat Genet, 2018. 50(5): p. 737–745.

2. Yang, J., et al., Common SNPs explain a large proportion of the heritability for human height. Nat Genet, 2010. 42(7): p. 565–9.

3. Bulik-Sullivan, B.K., et al., LD Score regression distinguishes confounding from polygenicity in genome-wide association studies. Nat Genet, 2015. 47(3): p. 291–5.

4. Timpson, N.J., et al., Genetic architecture: the shape of the genetic contribution to human traits and disease. Nat Rev Genet, 2018. 19(2): p. 110–124.

5. Holland, D., et al., Beyond SNP Heritability: Polygenicity and Discoverability of Phenotypes Estimated with a Univariate Gaussian Mixture Model. bioRxiv, 2019: p. 133132.

6. Frei, O., et al., Bivariate causal mixture model quantifies polygenic overlap between complex traits beyond genetic correlation. Nat Commun, 2019. 10(1): p. 2417.

7. Schaub, M.A., et al., Linking disease associations with regulatory information in the human genome. Genome Res, 2012. 22(9): p. 1748–59.

8. Schork, A.J., et al., All SNPs are not created equal: genome-wide association studies reveal a consistent pattern of enrichment among functionally annotated SNPs. PLoS Genet, 2013. 9(4): p. e1003449.

9. Finucane, H.K., et al., Partitioning heritability by functional annotation using genome-wide association summary statistics. Nat Genet, 2015. 47(11): p. 1228–35.

10. Zhu, X. and M. Stephens, Large-scale genome-wide enrichment analyses identify new trait-associated genes and pathways across 31 human phenotypes. Nat Commun, 2018. 9(1): p. 4361.

11. Zhang, Y., et al., Estimation of complex effect-size distributions using summary-level statistics from genome-wide association studies across 32 complex traits. Nat Genet, 2018. 50(9): p. 1318–1326.

12. Su, Z., J. Marchini, and P. Donnelly, HAPGEN2: simulation of multiple disease SNPs. Bioinformatics, 2011. 27(16): p. 2304–5.

13. Consortium, G.P., et al., A global reference for human genetic variation. Nature, 2015. 526(7571): p. 68–74.

14. Chang, C.C., et al., Second-generation PLINK: rising to the challenge of larger and richer datasets. Gigascience, 2015. 4: p. 7.

15. Karolchik, D., et al., The UCSC Table Browser data retrieval tool. Nucleic Acids Res, 2004. 32(Database issue): p. D493–6.

16. Frei, O. Precimed/SIMU GitHub page. 2016; Available from: https://github.com/precimed/simu.

17. Schizophrenia Working Group of the Psychiatric Genomics Consortium, Biological insights from 108 schizophrenia-associated genetic loci. Nature, 2014. 511(7510): p. 421–7.

18. Stahl, E.A., et al., Genome-wide association study identifies 30 loci associated with bipolar disorder. Nat Genet, 2019. 51(5): p. 793–803.

19. Demontis, D., et al., Discovery of the first genome-wide significant risk loci for attention deficit/hyperactivity disorder. Nat Genet, 2019. 51(1): p. 63–75.

20. Savage, J.E., et al., Genome-wide association meta-analysis in 269,867 individuals identifies new genetic and functional links to intelligence. Nat Genet, 2018. 50(7): p. 912–919.

21. Lee, J.J., et al., Gene discovery and polygenic prediction from a genome-wide association study of educational attainment in 1.1 million individuals. Nat Genet, 2018. 50(8): p. 1112–1121.

22. Mahajan, A., et al., Fine-mapping type 2 diabetes loci to single-variant resolution using high-density imputation and islet-specific epigenome maps. Nat Genet, 2018. 50(11): p. 1505–1513.

23. de Lange, K.M., et al., Genome-wide association study implicates immune activation of multiple integrin genes in inflammatory bowel disease. Nat Genet, 2017. 49(2): p. 256–261.

24. Willer, C.J., et al., Discovery and refinement of loci associated with lipid levels. Nat Genet, 2013. 45(11): p. 1274–1283.

25. Yengo, L., et al., Meta-analysis of genome-wide association studies for height and body mass index in approximately 700000 individuals of European ancestry. Hum Mol Genet, 2018. 27(20): p. 3641–3649.

26. Shungin, D., et al., New genetic loci link adipose and insulin biology to body fat distribution. Nature, 2015. 518(7538): p. 187–196.

27. LDSR data. Available from: https://data.broadinstitute.org/alkesgroup/LDSCORE/.

28. Minelli, C., et al., Importance of different types of prior knowledge in selecting genome-wide findings for follow-up. Genet Epidemiol, 2013. 37(2): p. 205–13.

29. Sveinbjornsson, G., et al., Weighting sequence variants based on their annotation increases power of whole-genome association studies. Nat Genet, 2016. 48(3): p. 314–7.

30. Flannick, J., et al., Exome sequencing of 20,791 cases of type 2 diabetes and 24,440 controls. Nature, 2019. 570(7759): p. 71-+.

31. Dewey, F.E., et al., Distribution and clinical impact of functional variants in 50,726 whole-exome sequences from the DiscovEHR Study. Science, 2016. 354(6319).

32. Speed, D., et al., Reevaluation of SNP heritability in complex human traits. Nat Genet, 2017. 49(7): p. 986–992.

33. Jones, E., E. Oliphant, and P. Peterson. SciPy: Open Source Scientific Tools for Python. 2001; Available from: https://www.scipy.org/.

34. Frei, O. and A.A. Shadrin. MiXeR GitHub page. 2016; Available from: https://github.com/precimed/mixer.

