## Supplementaty Tables and Figures for "Annotation-Informed Causal Mixture Modeling (AI-MiXeR) reveals phenotype-specific differences in polygenicity and effect size distribution across functional annotation categories"

### Supplementary material

#### Tables

| Phenotype | $\pi_{non-exonic}$ | $\pi_{exonic}$ | $\sigma^2_{non-exonic}$ | $\sigma^2_{exonic}$ | $\sigma^2_0$ | $N_{non-exonic}$ | $N_{exonic}$ | $h^2_{total}$ | $h^2_{non-exonic}$ | $h^2_{exonic}$ |
|---|---|---|---|---|---|---|---|---|---|---|
| SCZ | 3.11E-03<br>(2.81E-03,3.37E-03) | 2.24E-02<br>(1.86E-02,2.79E-02) | 5.30E-05<br>(4.87E-05,5.80E-05) | 1.04E-04<br>(8.27E-05,1.26E-04) | 1.13<br>(1.12,1.13) | 30610<br>(27709,33163) | 3384<br>(2817,4227) | 4.01E-01<br>(3.97E-01,4.08E-01) | 3.37E-01<br>(3.29E-01,3.42E-01) | 6.46E-02<br>(6.14E-02,6.92E-02) |
| BD | 2.77E-03<br>(2.44E-03,3.21E-03) | 1.83E-02<br>(1.54E-02,2.08E-02) | 4.05E-05<br>(3.53E-05,4.60E-05) | 1.17E-04<br>(1.01E-04,1.37E-04) | 1.06<br>(1.06,1.07) | 27322<br>(24031,31576) | 2767<br>(2336,3143) | 2.89E-01<br>(2.83E-01,2.94E-01) | 2.29E-01<br>(2.23E-01,2.36E-01) | 5.99E-02<br>(5.56E-02,6.35E-02) |
| ADHD | 1.47E-03<br>(1.30E-03,1.63E-03) | 3.86E-05<br>(1.06E-05,8.14E-05) | 5.69E-05<br>(5.29E-05,6.12E-05) | 1.65E-03<br>(8.72E-04,3.29E-03) | 1.11<br>(1.11,1.12) | 14426<br>(12769,16000) | 6<br>(2,12) | 1.72E-01<br>(1.63E-01,1.77E-01) | 1.70E-01<br>(1.62E-01,1.76E-01) | 1.71E-03<br>(4.28E-04,3.10E-03) |
| COG | 2.72E-03<br>(2.56E-03,2.88E-03) | 8.93E-03<br>(6.83E-03,1.11E-02) | 2.54E-05<br>(2.40E-05,2.67E-05) | 6.96E-05<br>(5.77E-05,8.93E-05) | 1.22<br>(1.21,1.23) | 26747<br>(25225,28359) | 1352<br>(1034,1676) | 1.58E-01<br>(1.56E-01,1.61E-01) | 1.41E-01<br>(1.39E-01,1.44E-01) | 1.73E-02<br>(1.59E-02,1.91E-02) |
| EA | 4.86E-03<br>(4.67E-03,5.20E-03) | 1.78E-03<br>(1.32E-03,2.17E-03) | 9.77E-06<br>(9.11E-06,1.01E-05) | 1.24E-04<br>(1.01E-04,1.51E-04) | 1.20<br>(1.19,1.20) | 47809<br>(45961,51168) | 270<br>(200,328) | 1.03E-01<br>(1.02E-01,1.04E-01) | 9.70E-02<br>(9.60E-02,9.83E-02) | 6.19E-03<br>(5.64E-03,7.11E-03) |
| T2D | 8.07E-05<br>(6.53E-05,9.30E-05) | 5.37E-02<br>(4.87E-02,6.25E-02) | 4.37E-04<br>(3.87E-04,5.52E-04) | 3.98E-05<br>(3.40E-05,4.41E-05) | 1.18<br>(1.18,1.19) | 795<br>(643,915) | 8134<br>(7368,9465) | 1.32E-01<br>(1.27E-01,1.36E-01) | 7.19E-02<br>(6.68E-02,7.50E-02) | 6.01E-02<br>(5.83E-02,6.29E-02) |
| IBD | 2.73E-05<br>(2.26E-05,3.28E-05) | 1.84E-02<br>(1.69E-02,1.99E-02) | 2.12E-03<br>(1.75E-03,2.47E-03) | 2.61E-04<br>(2.21E-04,2.95E-04) | 1.20<br>(1.19,1.20) | 269<br>(222,323) | 2788<br>(2555,3006) | 2.52E-01<br>(2.41E-01,2.62E-01) | 1.17E-01<br>(1.11E-01,1.27E-01) | 1.35E-01<br>(1.22E-01,1.43E-01) |
| LDL | 1.47E-04<br>(5.18E-05,2.06E-04) | 1.77E-03<br>(1.53E-03,2.52E-03) | 1.42E-04<br>(1.02E-04,3.07E-04) | 1.99E-03<br>(1.29E-03,2.35E-03) | 0.96<br>(0.95,0.98) | 1448<br>(510,2027) | 268<br>(232,381) | 1.39E-01<br>(1.24E-01,1.47E-01) | 4.12E-02<br>(3.26E-02,4.53E-02) | 9.82E-02<br>(9.11E-02,1.05E-01) |
| BMI | 3.64E-03<br>(3.50E-03,3.95E-03) | 4.05E-03<br>(3.39E-03,4.89E-03) | 2.33E-05<br>(2.16E-05,2.43E-05) | 1.67E-04<br>(1.42E-04,2.06E-04) | 1.61<br>(1.59,1.62) | 35834<br>(34414,38871) | 613<br>(513,741) | 1.93E-01<br>(1.91E-01,1.95E-01) | 1.74E-01<br>(1.72E-01,1.75E-01) | 1.90E-02<br>(1.79E-02,1.98E-02) |
| HEIGHT | 6.06E-04<br>(5.79E-04,6.27E-04) | 6.32E-02<br>(5.94E-02,6.96E-02) | 2.70E-04<br>(2.56E-04,2.83E-04) | 9.26E-05<br>(8.11E-05,9.94E-05) | 2.41<br>(2.38,2.44) | 5966<br>(5699,6169) | 9562<br>(8989,10532) | 4.99E-01<br>(4.90E-01,5.03E-01) | 3.34E-01<br>(3.27E-01,3.42E-01) | 1.65E-01<br>(1.59E-01,1.70E-01) |
| WHR | 4.17E-04<br>(3.76E-04,4.54E-04) | 6.97E-02<br>(4.68E-02,3.13E-01) | 4.92E-05<br>(4.56E-05,5.31E-05) | 8.01E-06<br>(1.56E-06,1.13E-05) | 0.94<br>(0.94,0.95) | 4107<br>(3697,4468) | 10556<br>(7082,47316) | 5.64E-02<br>(5.50E-02,5.79E-02) | 4.19E-02<br>(4.05E-02,4.36E-02) | 1.45E-02<br>(1.38E-02,1.52E-02) |

**Supplementary Table 1**. Estimates of proportion and number of non-null variants and their average effect sizes (mean values over 50 optimization runs are shown).
The numbers represent mean (min, max) values taken across 50 independent optimization runs with $10^6$ randomly selected variants used in the cost function for optimization.
$N_{exonic}$ = number of non-null variants in the exonic category = $\pi_{exonic} \times T_{exonic}$, where $T_{exonic}$ is the total number of variants in the exonic category. and $N_{non-exonic} = \pi_{non-exonic} \times T_{non-exonic}$, in the LDSC template used in our study $T_{non-exonic} \cong 70 * T_{exonic}$ and $T_{non-exonic} + T_{exonic} \cong 10^7$. Phenotypes: schizophrenia (SCZ), bipolar disorder (BD), attention deficit/hyperactivity disorder (ADHD), general cognitive ability (COG), educational attainment (EA), type 2 diabetes (T2D), inflammatory bowel disease (IBD), low-density lipoproteins (LDL), body mass index (BMI), height and waist-hip ratio (WHR).

| Phenotype | MiXeR | | | LDSC | | |
|---|---|---|---|---|---|---|
| | $h^2_{total}$ | Enrichment non-exonic | Enrichment exonic | $h^2_{total}$ | Enrichment non-exonic | Enrichment exonic |
| SCZ | 0.40<br>(4.73E-03,6.33E-03) | 0.85<br>(1.05E-02,8.92E-03) | 10.63<br>(5.80E-01,6.81E-01) | 0.37<br>(1.71E-02) | 0.91<br>(1.65E-02) | 7.64<br>(1.20E+00) |
| BD | 0.29<br>(5.58E-03,5.13E-03) | 0.80<br>(1.25E-02,1.23E-02) | 13.69<br>(7.97E-01,8.14E-01) | 0.27<br>(1.59E-02) | 0.86<br>(2.61E-02) | 11.19<br>(1.89E+00) |
| ADHD | 0.17<br>(8.81E-03,4.85E-03) | 1.01<br>(8.32E-03,7.64E-03) | 0.66<br>(4.97E-01,5.41E-01) | 0.19<br>(1.46E-02) | 1.03<br>(2.67E-02) | -1.11<br>(1.94E+00) |
| COG | 0.16<br>(2.23E-03,2.37E-03) | 0.90<br>(9.77E-03,9.86E-03) | 7.23<br>(6.41E-01,6.35E-01) | 0.16<br>(6.60E-03) | 0.94<br>(1.39E-02) | 5.69<br>(1.01E+00) |
| EA | 0.10<br>(1.13E-03,1.14E-03) | 0.95<br>(8.38E-03,5.31E-03) | 3.96<br>(3.46E-01,5.45E-01) | 0.10<br>(2.90E-03) | 0.97<br>(1.26E-02) | 2.97<br>(9.13E-01) |
| T2D | 0.13<br>(4.55E-03,4.01E-03) | 0.55<br>(2.06E-02,1.76E-02) | 30.06<br>(1.14E+00,1.34E+00) | 0.13<br>(9.50E-03) | 0.67<br>(4.25E-02) | 25.29<br>(3.09E+00) |
| IBD | 0.25<br>(1.15E-02,9.46E-03) | 0.47<br>(2.27E-02,2.66E-02) | 35.30<br>(1.73E+00,1.47E+00) | 0.23<br>(2.09E-02) | 0.55<br>(5.99E-02) | 33.57<br>(4.35E+00) |
| LDL | 0.14<br>(1.57E-02,7.84E-03) | 0.30<br>(3.28E-02,2.53E-02) | 46.51<br>(1.64E+00,2.13E+00) | 0.14<br>(1.73E-02) | 0.50<br>(9.13E-02) | 37.44<br>(6.63E+00) |
| BMI | 0.19<br>(1.68E-03,1.96E-03) | 0.92<br>(4.09E-03,5.72E-03) | 6.50<br>(3.72E-01,2.66E-01) | 0.19<br>(6.60E-03) | 0.92<br>(1.27E-02) | 6.56<br>(9.19E-01) |
| HEIGHT | 0.50<br>(8.81E-03,4.10E-03) | 0.68<br>(9.50E-03,1.29E-02) | 21.77<br>(8.36E-01,6.18E-01) | 0.47<br>(2.07E-02) | 0.77<br>(2.61E-02) | 17.37<br>(1.90E+00) |
| WHR | 0.06<br>(1.44E-03,1.48E-03) | 0.75<br>(1.20E-02,1.36E-02) | 16.96<br>(8.86E-01,7.83E-01) | 0.07<br>(5.50E-03) | 0.84<br>(3.10E-02) | 12.42<br>(2.25E+00) |

**Supplementary Table 2**. Comparison of partitioned heritability estimates from AI-MiXeR and LDSC.
For AI-MiXeR, confidence intervals are presented as (mean – min, max - mean) where mean, min and max are taken across 50 runs of cost function optimization with $10^6$ randomly selected variants. For LDSC, standard errors are given in brackets.

**Supplementary Table 3**. Results for all 810 simulation scenarios (in separate Excel file).

| Trait | Year of publication | Sample size (total or cases/controls) |
|---|---|---|
| Schizophrenia (SCZ), 49 European sub-studies | 2014 | 33640/43456 |
| Bipolar disorder (BD) | 2019 | 20352/31358 |
| Attention deficit/hyperactivity disorder (ADHD) | 2019 | 19099/34194 |
| General cognitive ability (COG) | 2018 | 269867 |
| Educational attainment (EA) | 2018 | 766345 |
| Type 2 diabetes (T2D) | 2018 | 74124/824006 |
| Inflammatory bowel disease (IBD) | 2017 | 25042/34915 |
| Low-density lipoproteins (LDL) | 2013 | 188577 |
| Body mass index (BMI) | 2018 | 795640 |
| Height | 2018 | 709706 |
| Waist-hip ratio (WHR) | 2015 | 224459 |

**Supplementary Table 4**. Details of GWAS on 11 phenotypes analyzed in the study.

#### Figures

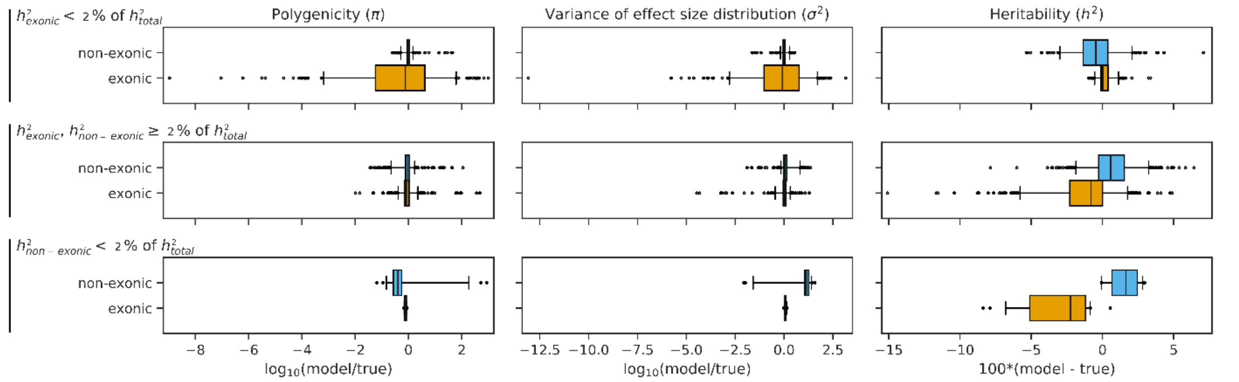


**Supplementary Figure 1.** Performance of the model on synthetic GWAS data.
Three different scenarios: $h^2_{exonic}$ < 2% of $h^2_{total}$, 300 tests (top row), both $h^2_{exonic}$ and $h^2_{non-exonic}$ >= 2% of $h^2_{total}$, 480 tests (middle row), and $h^2_{non-exonic}$ < 2% $h^2_{total}$, 30 tests (bottom row).
The color coding defines the functional annotation category (blue: non-exonic, orange: exonic).
For polygenicity (first column) and discoverability (second column), the X axis shows by how many orders of magnitudes the modeled parameter is greater than corresponding real parameter: $\log_{10}$(model/true). For heritability, the X axis shows the difference between modeled and true heritabilties (measured in percents, i.e. $0 \leq h^2 \leq 100$). For all subplots, closer to 0corresponds to better performance of the model.
Boxes include 50% of all tests, whiskers show 95% confidence intervals, dots represent outliers.

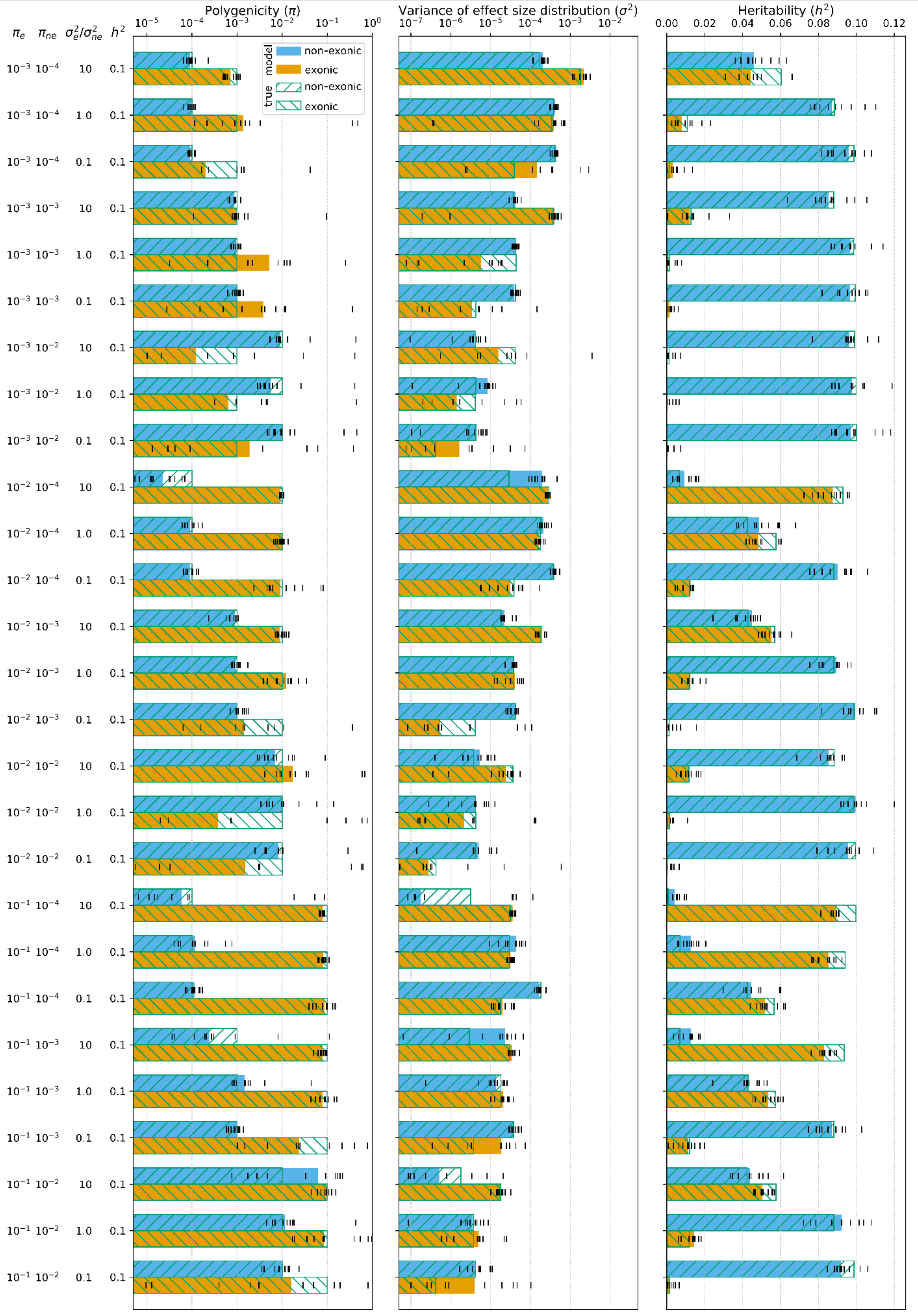


**Supplementary Figure 2.** Simulations with $h^2_{total}$ = $h^2_{exonic}$+$h^2_{non-exonic}$ = 0.1 and all combinations of $\pi_{exonic}$ = $10^{-1}$, $10^{-2}$, $10^{-3}$; $\pi_{non-exonic}$ = $10^{-2}$, $10^{-3}$, $10^{-4}$ and $\sigma^2_{exonic}/\sigma^2_{non-exonic}$ = 0.1, 1.0, 10.0.

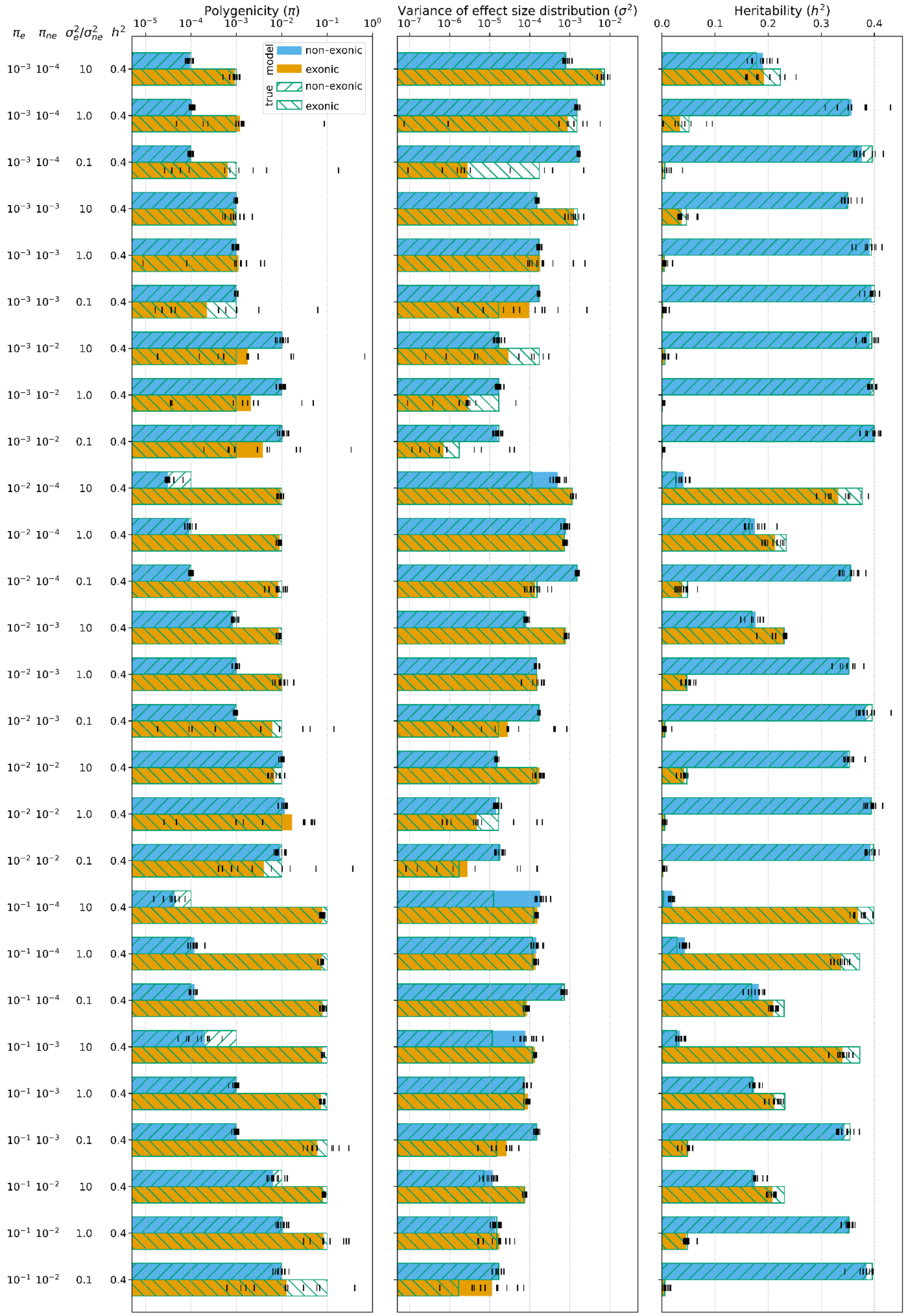


**Supplementary Figure 3.** Simulations with $h^2_{total} = h^2_{exonic} + h^2_{non-exonic}$ = 0.4 and all combinations of $\pi_{exonic}$ = $10^{-1}$, $10^{-2}$, $10^{-3}$; $\pi_{non-exonic}$ = $10^{-2}$, $10^{-3}$, $10^{-4}$ and $\sigma^2_{exonic}/\sigma^2_{non-exonic}$ = 0.1, 1.0, 10.0.

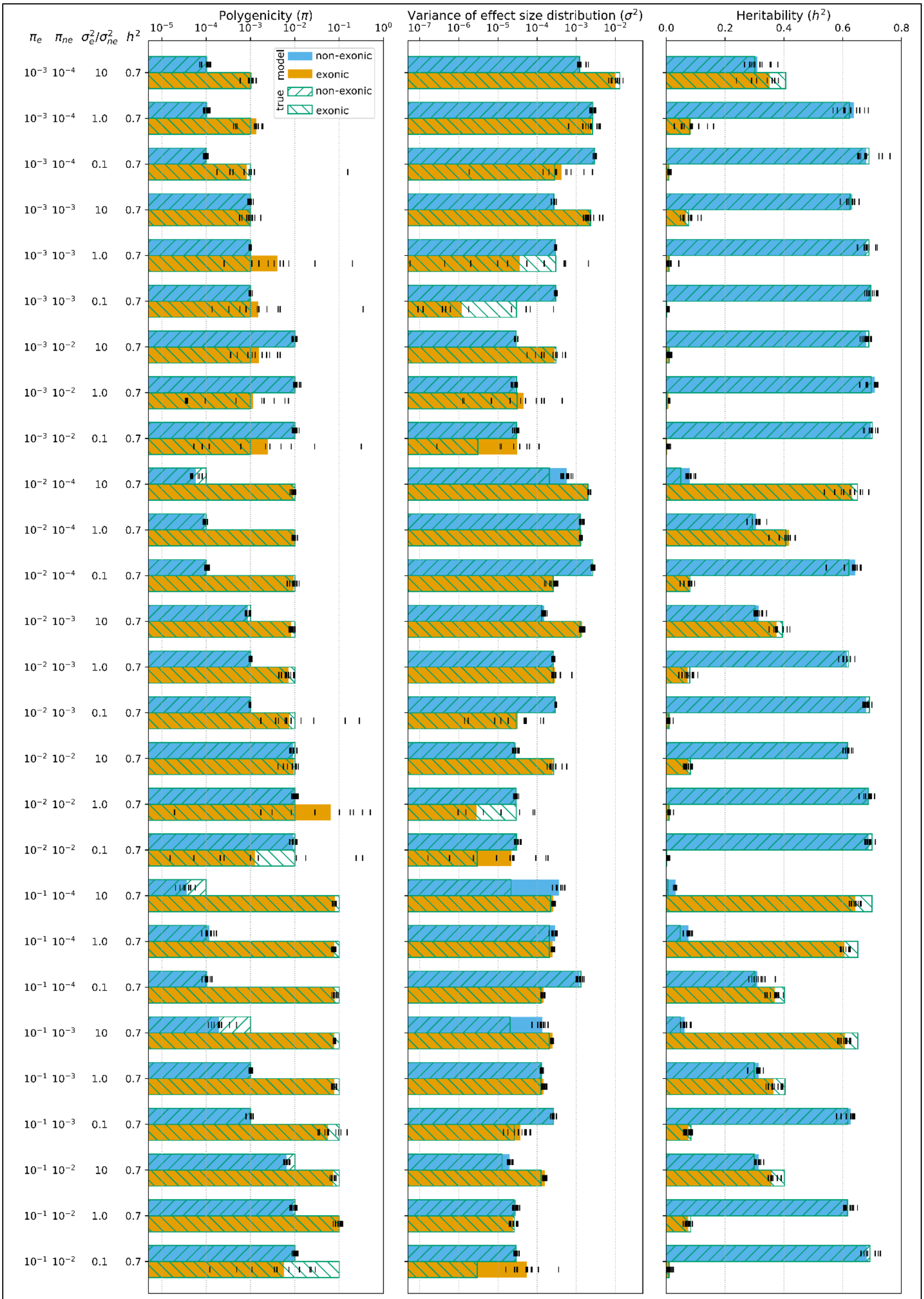


**Supplementary Figure 4.** Simulations with $h^2_{total}$ = $h^2_{exonic}$+$h^2_{non-exonic}$ = 0.7 and all combinations of $\pi_{exonic}$ = $10^{-1}$, $10^{-2}$, $10^{-3}$; $\pi_{non-exonic}$ = $10^{-2}$, $10^{-3}$, $10^{-4}$ and $\sigma^2_{exonic}/\sigma^2_{non-exonic}$ = 0.1, 1.0, 10.0.
